# Adaptation of an herbivorous arthropod to green tea plants by overcoming catechin defenses

**DOI:** 10.64898/2025.12.13.694159

**Authors:** Naoki Takeda, Brendan Abiskaroon, Ricardo Hernandez Arriaza, Ryutaro Murakami, Shogo Sasaki, Masanobu Yamamoto, Vladimir Zhurov, Vojislava Grbić, Maksymilian Chruszcz, Takeshi Suzuki

## Abstract

Green tea catechins are known antioxidants that benefit human health and protect tea plants from biotic stressors. However, some herbivores can counteract catechin defenses and can use tea plants as a host. Among herbivorous mites, an extreme generalist *Tetranychus urticae* has not been reported as a tea pest. Instead, *T. kanzawai*, another generalist, has some populations that thrive on tea plants. Here, we investigated the mechanism of the adaptation of these mites to tea plants. Comparative study of the intra- and inter-specific variations in mite performances uncovered differences in their behavioral and xenobiotic responsiveness to green tea catechins. We showed that green tea catechins exert complex defensive roles. They were repellent and toxic to *T. urticae* and tea non-adapted *T. kanzawai* mites. In addition, they had an antifeedant effect on tea non-adapted *T. kanzawai* mites. Matching the catechin structure, we identified an intradiol ring-cleavage dioxygenase *DOG15*, a gene horizontally transferred from fungi, as one required for the adaptation of *T. kanzawai* mites to tea plants. The *DOG15* gene has an enhanced inducible expression in tea-adapted *T. kanzawai* mites, with mRNA and protein levels up to 31.6 and 12.1 times higher than in *T. urticae* mites fed on tea plants. Furthermore, we identified two amino acid substitutions in DOG15 between *Tetranychus* species leading to the increased efficacy of the *T. kanzawai* encoded enzyme toward cleavage of green tea catechins. Thus, we showed that mite adaptation to tea plants occurred in a two-step process. The amino acid substitutions in DOG15 predispose *T. kanzawai* but not *T. urticae* for the adaptation to tea plants. Further increased expression of modified DOG15 enables *T. kanzawai* mites to efficiently detoxify green tea catechins, leading to intra- and inter-specific differences in mites’ ability to use tea plants as a host. Our findings reveal how a horizontally transferred gene can be co-opted through structural and regulatory changes to overcome plant chemical defenses, with implications for herbivore host adaptation and tea pest management.

## Introduction

Green tea is made from unfermented leaves of the plant *Camellia sinensis* (L.) O. Kuntze. Unlike black tea, which is fully oxidized during the fermentation process, green tea contains higher levels of condensed flavonoids called catechins, antioxidants shown to have a variety of health benefits, including anticancer and antimicrobial activities (Cabrera et al. 2006). Green tea catechins comprise 2.4% to 23.8% of the dry mass of tea leaves (Baptista et al., 2014; Deka et al., 2021). The major catechins in tea leaves are (−)-epigallocatechin gallate (EGCg), (−)-epigallocatechin (EGC), (−)-epicatechin gallate (ECg), and (−)-epicatechin (EC). While these catechins have health benefits for humans, they are used by the tea plants as defensive compounds to protect them against pest fungi and herbivores (Ullah et al., 2017; Li et al., 2022). At present, it is unclear why some herbivores can survive and reproduce on catechin-rich tea plants and others cannot. In mammals, green tea catechins are metabolized in the liver and small intestine through processes such as *O*-methylation, sulfation, and glucuronidation. These metabolic pathways facilitate the conversion of catechins to more water-soluble forms, which enhance their excretion via urine and bile (Chu and Pang, 2018).

The herbivorous mites are an excellent model to address the differential ability of herbivores to adapt to tea plants. On one hand, the extremely polyphagous *Tetranychus urticae* Koch (Trombidiformes: Tetranychidae), the two-spotted spider mite (TSSM), that feeds on more than 1,100 species including economically important agricultural and horticultural crops (Migeon and Dorkeld, 2025) has not been reported on tea plants. However, the closely related Kanzawa spider mite (KSM), *T. kanzawai* Kishida, is one of the most important pests of tea plants (Hazarika et al., 2009). Intraspecific variation divides KSM into adapted and non-adapted populations to tea plants (Gomi and Gotoh, 1996). The intra- and inter-specific variation in the ability of mites to infest tea plants may reflect differences in how they avoid or detoxify green tea catechins, the major defense compounds.

Unlike humans, spider mites have 17 genes encoding intradiol ring-cleavage dioxygenases (DOGs) acquired through a single horizontal gene transfer (HGT) from fungi (Grbić et al., 2011; Dermauw et al., 2013). In bacteria and fungi, DOGs are characterized by their ability to catalyze the oxidative cleavage of aromatic rings in catecholic compounds like catechins between adjacent hydroxyl groups (Guzik et al., 2013). Spider mite DOGs can also directly use catechins as a substrate in *in vitro* reactions (Njiru et al. 2022), raising the possibility that they may carry the more efficient metabolic transformation of catechins than the multi-step pathways observed in mammals. However, despite the enzymatic capability, the adaptive significance of the DOG genes in relation to mite host adaptation has not been reported. In the present study, we aimed to determine the molecular basis by which KSM, but not TSSM, adapts to the catechin defenses of tea plants. We hypothesized that this adaptation involves both reduced chemosensory avoidance of catechins and enhanced enzymatic detoxification. To characterize these components and identify the genes underlying adaptation, we exploited intra- and inter-specific variation in spider mites’ host utilization of tea plants, combining behavioral and toxicological assays with comparative transcriptomics and proteomics. We then functionally validated candidate genes by RNAi-mediated gene silencing and recombinant enzyme assays. These analyses revealed reduced chemosensory avoidance in tea-adapted KSM and identified the functional role of DOG15 enzyme in mite adaptation to catechins. Our findings provide the first empirical evidence linking DOG enzymatic function to host plant specialization and reveal the molecular basis by which KSM has emerged as a tea pest.

## Results

### Effects of tea leaves and catechins on mite performance

To investigate the mechanisms underlying tea adaptation, we used a tea-adapted KSM population collected from a tea farm, along with tea non-adapted KSM and TSSM that do not naturally utilize tea as a host. We compared the feeding activity, survival, and fecundity of three populations on leaves of tea and kidney bean (*Phaseolus vulgaris* L.), a generally preferred host of *Tetranychus* mites, to assess whether tea adaptation confers measurable fitness changes. Tea leaf discs with tea-adapted KSM formed dark color patterns due to mite feeding, while those with non-adapted KSM and TSSM showed little or no such leaf damage (Figures 1a, b). Survival and fecundity on tea leaves were significantly higher in tea-adapted KSM than in non-adapted KSM and TSSM (Figures 1c, d). After 10 days, almost 90% of tea-adapted KSM survived, compared with about 5% of non-adapted KSM and 33% of TSSM. Tea-adapted KSM laid up to about 2 eggs/surviving female daily, whereas the other two populations laid almost no eggs. When tea-adapted KSM were transferred onto the leaves of kidney bean, their fecundity was significantly higher than that on tea leaves and there was no significant reduction in their survival (Supplemental Figures 1-1a, b). In TSSM, the survival and fecundity were significantly higher on kidney bean leaves than on tea leaves.

**Figure 1.**
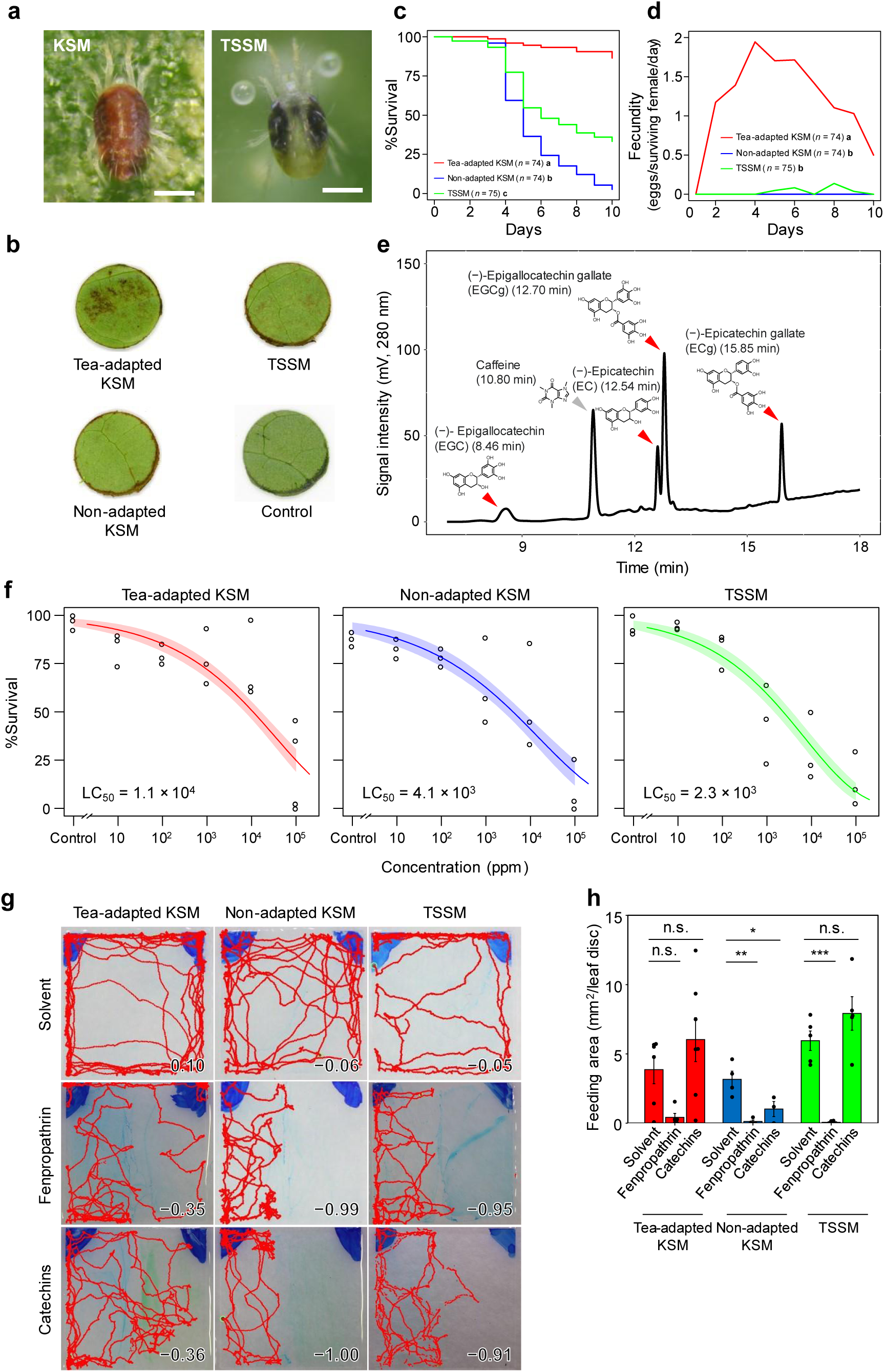
Mite performance on tea leaves and sensitivity to green tea catechins. (a) Adult females of the Kanzawa spider mite (KSM) *Tetranychus kanzawai* and the two-spotted spider mite (TSSM) *Tetranychus urticae*. Scale bars indicate 200 µm. (b) Feeding damage on leaf discs (10 mm in diameter) of the tea plant *Camellia sinensis* by tea-adapted KSM, non-adapted KSM, and TSSM. Each leaf disc was fed by 15 adult females at 25°C for 1 day. Control indicates a mite-free leaf disc. (c) Survival and (d) fecundity of adult females of tea-adapted KSM (*n* = 74), non-adapted KSM (*n* = 74), and TSSM (*n* = 75) on leaves of tea seedlings at 25 °C for 10 days. In each population, mites that had grown to the teleiochrysalid stage on leaves of kidney bean *Phaseolus vulgaris* and molted to adults within 2 h were transferred to a single tea leaf and the numbers of survivors and eggs-laid were counted daily. Mites that escaped from the leaves during the experiment were excluded from the data. Data were collected from 5 independent experimental runs (*n* = 14 to 15 per run) using different single leaves. Different letters indicate a significant difference (*p* < 0.001) in the survival and fecundity among mite populations by log-rank test with Bonferroni correction and Tukey’s HSD test, respectively. (e) HPLC chromatogram of green tea catechins in leaves of tea seedlings used in the mite performance test. Red arrowheads and a gray arrowhead indicate green tea catechins and caffeine, respectively. The numbers in parentheses following the compound names indicate the retention time for each peak. (f) Dose–response relationships between catechin levels and survivorship of tea-adapted KSM, non-adapted KSM, and TSSM, exposed to different concentrations (ppm, w/v) of an aqueous solution of a catechin mixture at 25 °C for 1 day. The catechin mixture consisted of (−)-epigallocatechin gallate (EGCg), (−)-epigallocatechin (EGC), (−)-epicatechin gallate (ECg), and (−)-epicatechin (EC). The number of survivors was counted at 3 days after recovery from soaking in the catechin solution. Data were collected from 3 to 5 independent experimental runs (*n* = 25 to 45 per run) and are presented with a regression line and a 95% confidence band. (g) Chemo-orientation behavior to green tea catechins in tea-adapted KSM, non-adapted KSM, and TSSM. The red curves indicate the trajectory of the mite’s locomotion for 10 min on a glass plate (9.0 × 9.0 mm), one half (4.5 × 9.0 mm) coated with the 10% (w/v) catechin mixture, 10 mM fenpropathrin knwon as a mite repellent substance (positive control), or their solvent consisting of 0.1% (w/v) brilliant blue FCF and 0.1% (v/v) Tween 20 in methanol (negative control), and the other half (4.5 × 9.0 mm) with the solvent. The numerical value at the bottom right of each image indicates a preference index (PI; see Supplemental Figure 1-2 for the assay procedure and index definition). (h) Feeding preference for green tea catechins in tea-adapted KSM, non-adapted KSM, and TSSM. Kidney bean leaf discs (10 mm in diameter) were coated with the 10% (w/v) catechin mixture, 10 mM fenpropathrin (positive control) in methanol, or their solvent methanol (negative control) and were then covered with stretched Parafilm to prevent direct contact of mites’ legs with the coated compounds. Adult females from each population were transferred onto leaf discs (15 mites/disc) and allowed to feed by penetrating the stretched Parafilm with their stylets at 25 °C for 24 h. Data are presented as the mean ± SE and collected from 3 to 7 independent experimental runs and compared using Dunnett’s test.

Tea leaves are rich in catechins, a group of condensed flavonoids that function as plant defense compounds against herbivores. To determine whether observed performance differences were due to differential tolerance to catechins, we analyzed catechin content in tea leaves and conducted their toxicity assays on three mite populations. High-performance liquid chromatography (HPLC) analysis revealed that leaves of the tea seedlings used in this study contained the four major catechins EGCg, EGC, ECg, and EC (Figure 1e). We then tested the catechin toxicity to mites by soaking them in the mixture of these four detected catechins. The median lethal concentration (LC_50_) values and the 95% confidence limits were 1.1 × 10^4^ ppm (6.3 × 10^3^ to 1.5 × 10^4^ ppm) for tea-adapted KSM, 4.1 × 10^3^ ppm (2.0 × 10^3^ to 6.1 × 10^3^ ppm) for non-adapted KSM, and 2.3 × 10^3^ ppm (1.3 × 10^3^ to 3.4 × 10^3^ ppm) for TSSM (Figure 1f). This indicates that tea-adapted KSM are less sensitive to green tea catechins than the other two populations.

Plant defense compounds can affect herbivores not only through toxicity but also by inducing avoidance behaviors, which is one of the most effective defense strategies for plants. Since catechins are non-volatile compounds, we evaluated their potential roles in contact-mediated chemical defenses by assessing both irritancy and antifeedancy. To assess the chemo-orientation behavior of mites to green tea catechins, adult females of tea-adapted KSM, non-adapted KSM, and TSSM were transferred onto 9.0 × 9.0 mm glass plates (1 mite/plate), on which one half (9.0 × 4.5 mm) and the other half (9.0 × 4.5 mm) were coated with the catechin mixture or fenpropathrin (positive control; Penman et al., 1986) and the solvent, respectively (Supplemental Figure 1-2a). While tea-adapted KSM did not avoid catechins nor fenpropathrin, non-adapted KSM and TSSM clearly showed contact irritancy to avoid them (Figure 1g, Supplemental Figure 1-2b, and Supplemental Movies 1 to 9). To quantify mite feeding activity, adult females of tea-adapted KSM, non-adapted KSM, and TSSM were transferred onto kidney bean leaf discs (1 mite/disc) coated with green tea catechins, fenpropathrin, or methanol. The leaf discs were covered with stretched Parafilm to allow the mite stylets to penetrate the film and reach the underlying leaf tissue for feeding. This setup prevented the mite legs from contacting the leaf surface directly, thus preventing chemosensory detection that would induce contact irritancy. While tea-adapted KSM and TSSM showed feeding on catechin-coated leaf discs, no feeding was observed in non-adapted KSM (Figure 1h). These results show that green tea catechins act as behavioral defenses, deterring mites through both irritancy and antifeedancy, to which tea-adapted KSM are largely insensitive.

**Figure 2.**
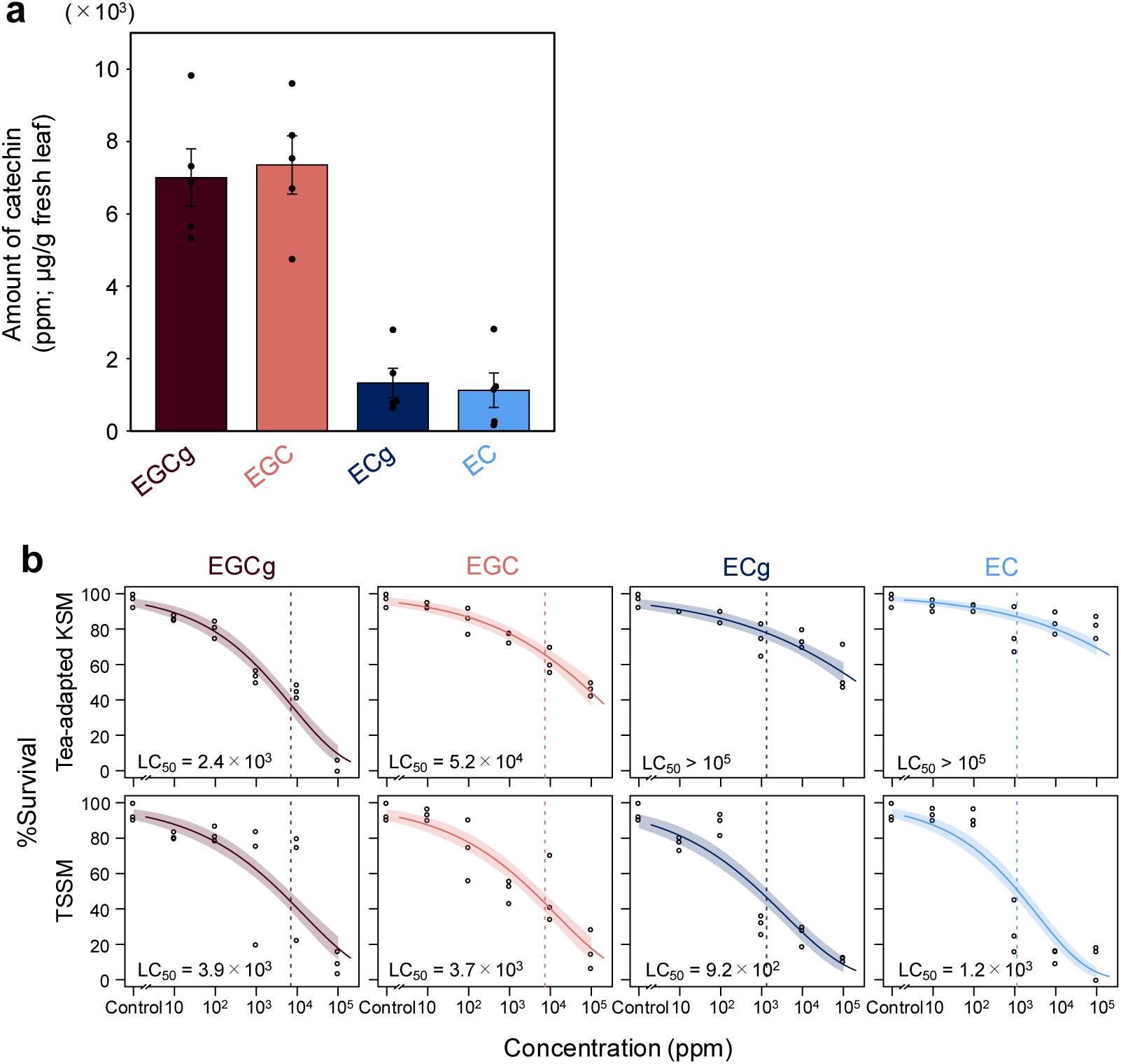
Green tea catechins in tea leaves and their toxicity to spider mites. (a) The amount of green tea catechins in fresh tea leaves, quantified by HPLC for (−)-epigallocatechin gallate (EGCg), (−)-epigallocatechin (EGC), (−)-epicatechin gallate (ECg), and (−)-epicatechin (EC). Data were collected from 5 independent experimental runs and are presented as the mean ± SE. (b) Catechin dose–response curves of survival of tea-adapted KSM and TSSM exposed to different concentrations (ppm, w/v) of an aqueous solution of EGCg, EGC, ECg, or EC at 25 °C for 24 h. Data were collected from 3 to 5 independent experimental runs (*n* = 28 to 51 per run) and are presented with a regression curve and a 95% confidence interval. The dotted lines indicate the average amounts of each catechin contained in fresh tea leaves.

### Catechin content in tea leaves and its toxicity to spider mites

To determine which catechins contribute to the interspecific differences in mite performance between KSM and TSSM, we first quantified four major catechins detected in fresh tea leaves. HPLC analysis revealed that the epigalloylated catechins EGCg and EGC were present at comparable concentrations (∼7 × 10^3^ µg/g fresh leaf each), which were approximately seven times higher than those of the non-galloylated catechins ECg and EC (∼1 × 10^3^ µg/g fresh leaf each) (Figure 2a). Based on approximately 80% moisture in fresh tea leaves (Wu et al., 2022), these values correspond to total catechin contents of approximately 84 mg/g (=8.4%) of the dry mass. This value was comparable to 2.4% to 23.8% of the dry mass of tea leaves (Baptista et al., 2014; Deka et al., 2021). No catechins were detected in fresh kidney bean leaves. The miticidal effects of EGCg, EGC, ECg, and EC were then evaluated. Newly molted adult females of tea-adapted KSM and TSSM were soaked in an aqueous solution of each catechin at 0 (control), 10, 10^2^, 10^3^, 10^4^, and 10^5^ ppm for 24 h. Based on the LC_50_ values (Supplemental Table 2-1), the sensitivity of tea-adapted KSM relative to TSSM was comparable for EGCg and approximately 14-fold lower for EGC. For ECg and EC, the LC_50_ values of tea-adapted KSM exceeded the highest concentration tested (10^5^ ppm), indicating markedly lower sensitivity than TSSM (Figure 2b).

### Inducible expression of *TkDOG15* is required for KSM adaptation to tea

To identify genes underlying host adaptation in tea-adapted KSM, we compared its transcriptome and proteome with those of non-adapted KSM and TSSM. We also compared mite feeding on tea versus bean leaves to distinguish between constitutive and feeding-inducible expression. Principal component analysis of the expression data of both mRNA (Figure 3a) and protein (Figure 3b) partitioned samples into the groups of KSM and TSSM along PC1. In addition, for both data sets, samples were partitioned into the groups of mites feeding on preferred and nonpreferred host plants along PC2. While TSSM and tea non-adapted KSM showed differential responses when feeding on bean versus tea, the tea-adapted KSM showed similar patterns when fed on leaves of these plant hosts. Comparison of inter-specific responses between the tea-adapted KSM and TSSM when fed on tea leaves, detected 1146 mRNAs and 601 proteins that were highly enriched (log2 fold change ≥ 1 with adjusted *p*-value < 0.05 for mRNA and *p*-value < 0.05 for protein; see Materials and Methods). To identify genes associated with KSM adaptation to tea plants, we compared intra-specific responses between tea-adapted and non-adapted KSM strains when fed on tea leaves. This contrast identified 756 mRNAs and 399 proteins. Finally, to identify the effect of plant host on tea-adapted KSM, we compared the transcriptome and proteome of these mites when fed on tea and bean leaves. This analysis identified 165 mRNAs and 329 proteins that were highly enriched. From these transcriptome and proteome comparisons, 32 mRNAs and 29 proteins were common, identifying genes and proteins that are specifically enriched in tea-adapted KSM feeding on tea (Supplemental Figures 3-1a, b and Supplemental Tables 3-1, 3-2). Among these, the gene encoding TkDOG15, the KSM ortholog of *tetur20g01790*, was the only gene/protein that was shared between enriched mRNAs and proteins (Figure 3c). In tea-adapted KSM fed on tea leaves, the mRNA and protein levels of the *TkDOG15* gene were 19.7 and 4.6 times higher than those fed on kidney bean leaves, 12.1 and 3.5 times higher than those in non-adapted KSM fed on tea leaves, and 31.6 and 12.1 times higher than those in TSSM fed on tea leaves, respectively (Figures 3d, e). To test if *TkDOG15* is required for KSM adaptation to tea, we silenced its expression using RNAi. Feeding tea-adapted KSM with dsRNA that specifically targets the *TkDOG15* gene decreased its expression by ∼75% and resulted in a significant reduction of their survival on tea leaves (Figures 3f, g), demonstrating that high expression of *TkDOG15* is required for KSM adaptation to tea.

**Figure 3.**
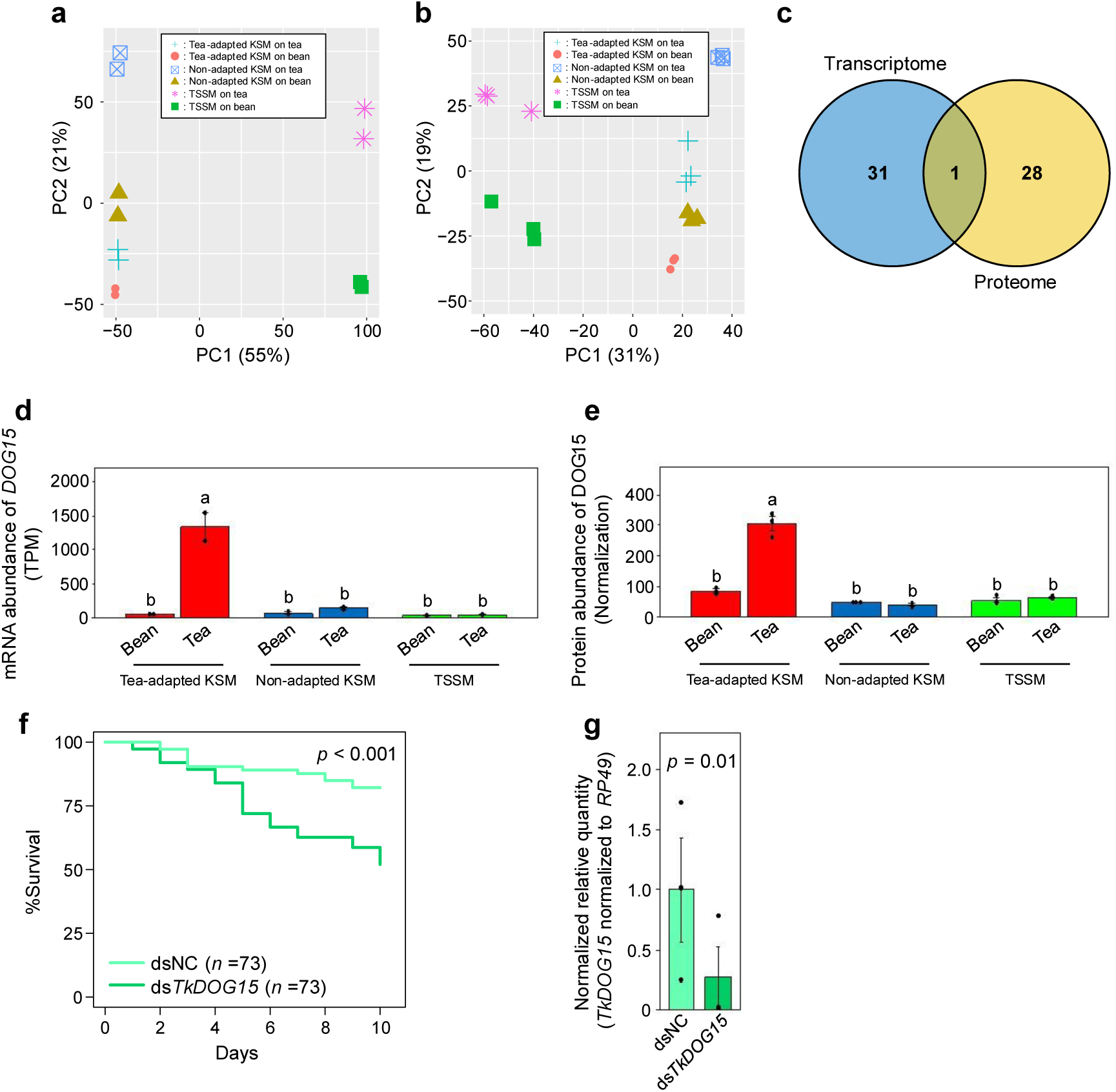
Gene expression patterns at the mRNA and protein levels in spider mites in response to consumption of tea or bean leaves revealed a specific high expression of the horizontally transferred gene *TkDOG15* in tea-adapted KSM fed on tea leaves. Principal component analysis of (a) transcriptome and (b) proteome of tea-adapted KSM, non-adapted KSM, and TSSM fed on tea or kidney bean leaves at 25 °C for 24 h. Data were collected from 2 (transcriptome) and 3 (proteome) independent experimental runs (*n* = 150 per run). (c) Comparative transcriptomics and proteomics revealed 32 mRNAs and 29 proteins that were highly expressed in tea-adapted KSM fed on tea leaves compared to tea-adapted KSM fed on bean leaves, non-adapted KSM fed on tea leaves, and TSSM fed on tea leaves. Among them, *TkDOG15* was the only gene that was commonly upregulated at both mRNA and protein levels in tea-adapted KSM fed on tea leaves. For the *DOG15* gene, (d) mRNA and (e) protein expression levels in the three populations fed on tea and kidney bean leaves. Expression levels were analyzed from the transcriptome and proteome data. Data are presented as the mean ± SE and different letters indicate a significant difference between treatments by Tukey’s HSD test. (f) Survival of tea-adapted KSM on tea leaves for 10 days after RNAi-mediated silencing of the *TkDOG15* gene. Mites that escaped from the leaf during the experiment were excluded from the data. Data were collected from 5 independent experimental runs (*n* = 15 per run). Kaplan-Meier survival curves were compared using the log-rank test. (g) Expression of the *TkDOG15* gene in adult females of tea-adapted KSM relative to that of the *PR49* gene at 4 days after the RNAi treatment with ds*TkDOG15* and dsNC. Data are presented as the mean ± SE and compared using Student’s *t*-test.

### TkDOG15 cleaves catechins more efficiently than TuDOG15

Our transcriptome and proteome analyses revealed that DOG15 was exclusively upregulated in tea-adapted KSM when feeding on tea leaves, compared to non-adapted KSM and TSSM. To assess whether DOG15 function in tea-adapted KSM differs not only quantitatively but also qualitatively, we examined amino acid sequence variations between TkDOG15 and TuDOG15, and their enzymatic activities against catechins as substrates. DOG15 has a conserved active site consisting of two histidine (H208 and H210) and two tyrosine (Y111 and Y202) residues that coordinate Fe^3+^ (Figure 4a, b, c). Amino acid sequences of DOG15 are identical between tea-adapted KSM and non-adapted KSM, whereas there are two amino acid substitutions (Q127A and T203A) in DOG15 between KSM (TkDOG15) and TSSM (TuDOG15) (Figure 4a). T203A is adjacent to Y202, one of the residues within the active site. To test if these polymorphisms affect the enzymatic activity of DOG15, we expressed TkDOG15 and TuDOG15 in *E. coli* and tested their abilities to use tea catechins as a substrate. The activity of TkDOG15 was significantly higher than that of TuDOG15 against EGCg, EGC, and EC (Supplemental Table 4-1). When the enzymatic activity was normalized to the abundance of DOG15 expressed in mites fed on tea plants (Figure 3e), TkDOG15 showed distinctly higher activity relative to TuDOG15 for all four green tea catechins (Figure 4c).

**Figure 4.**
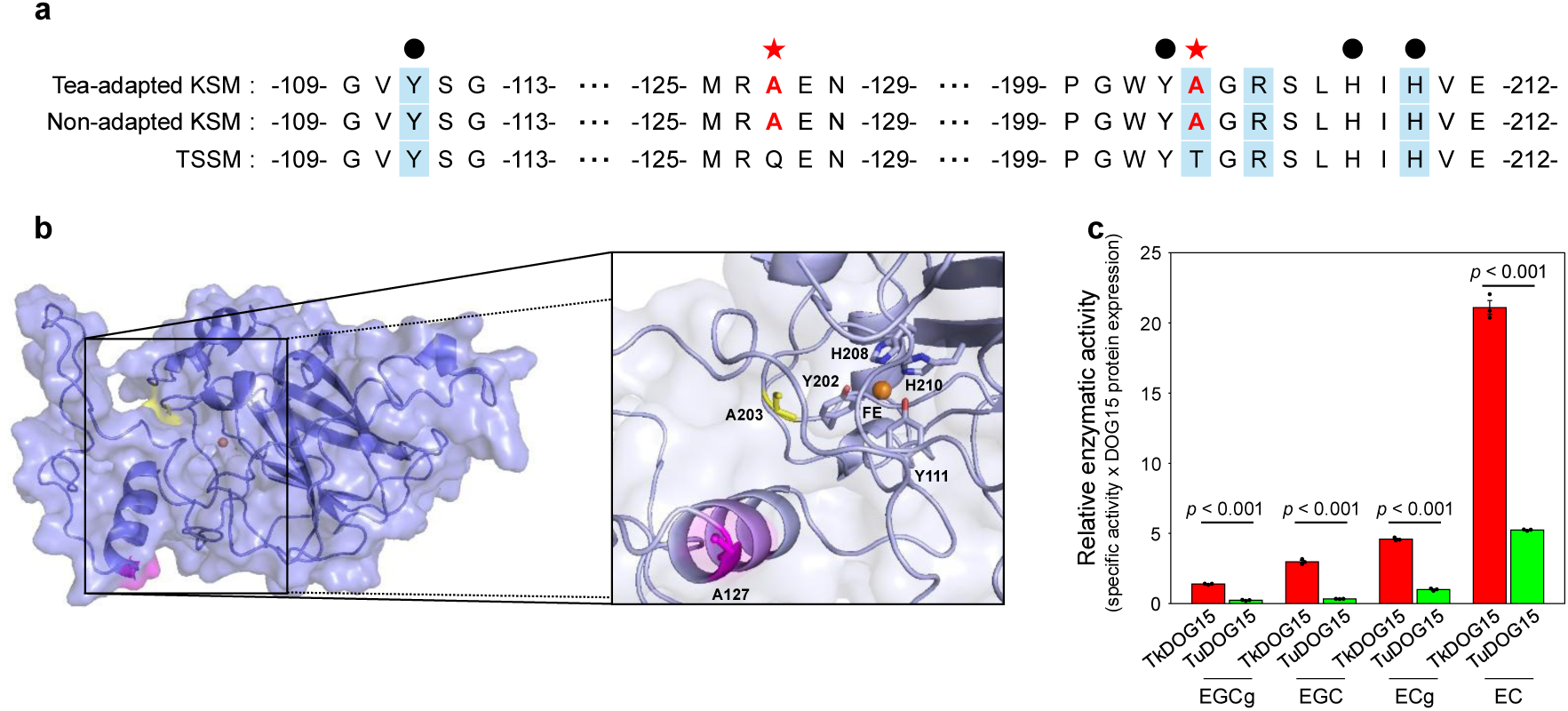
Comparisons of DOG15 structure and enzymatic activity between KSM and TSSM. (a) Amino acid sequences of DOG15 in tea-adapted KSM, non-adapted KSM, and TSSM, aligned using the MUSCLE algorithm. The amino acid sequences of tea-adapted and non-adapted KSM were completely matched, but amino acid substitutions were confirmed at Q127A and T203A (red star) between KSM and TSSM. The active sites based on the two tyrosine (Y111 and Y202) and histidine (H208 and H210) residues were matched in KSM and TSSM (black dot). (b) AlphaFold2 model of TkDOG15 showing differing residues between TkDOG15 (A127 in pink; A203 in yellow) and TuDOG15 (Q127 and T203; not shown). Two tyrosine and two histidine residues coordinating iron cofactor (orange) are displayed in the active site. (c) Relative enzymatic activity to green tea catechins in TkDOG15 and TuDOG15. Relative enzymatic activity was calculated by multiplying the specific activity of DOG15 for each catechin by the abundance of DOG15 in tea-adapted KSM and TSSM on tea plants (Figure 3e), respectively. Data are presented as the mean ± SD and compared using Student’s *t*-test.

## Discussion

Because plants have limited mobility, they protect themselves by producing a variety of defensive compounds that disrupt behavior and/or are toxic to herbivores (Koul, 2008). We found that green tea catechins affect mites through contact irritancy (Figure 1g), antifeedancy (Figure 1h) and toxicity (Figure 1f). Although there are no reports of contact irritancy of green tea catechins in arthropods, antifeedancy of EGCg, one of the most abundant green tea catechins (Figure 2a), was reported in the cotton aphid, *Aphis gossypii* Glover (Zhao et al., 2021). Spider mites possess chemosensory organs on their mouthparts, pedipalps, and legs I and II (Tateishi, 1988; Sakunwarin et al., 2004). Catechins, localized in mesophyll cell vacuoles (Suzuki et al., 2003), are expected to be released during stylet penetration (Bensoussan et al., 2016, 2018; Arai et al., 2025), potentially triggering contact irritancy and antifeedant responses. Nevertheless, tea-adapted KSM overcome these behavioral barriers. Spider mites possess an exceptionally large repertoire of chemosensory receptors (Ngoc et al., 2016; Chen et al., 2023), among which some likely function as catechin receptors that mediate the observed behavioral responses in non-adapted KSM and TSSM. We hypothesize that the insensitivity of tea-adapted KSM to the repellency of tea catechins arises either from loss-of-function mutations in catechin-responsive receptor(s) or from rapid detoxification of catechins within the feeding cell before they reach the leaf surface. Identifying the responsible receptors and tracing the fate of catechins within the feeding cell remain to be addressed in future studies.

Tea-adapted KSM counteracted the toxicity of green tea catechins with TkDOG15 (Figure 3c-e), a catechol dioxygenase that oxidatively cleaves catechol or pyrogallol rings (Figure 4a-d). The metabolism of green tea catechins, involving *O*-methylation, sulfation, and glucuronidation, has only been reported in mammals, where it increases water solubility and enhances excretion via urine and bile (Chu and Pang, 2018). The DOG-mediated system for detoxifying catechins in spider mites is simpler than the mammalian system. Herbivorous arthropods demonstrate an evolutionary trend toward utilizing plants as dietary resources through optimization of detoxification mechanisms that counteract toxic secondary metabolites, a key component of plant chemical defense systems (Schoonhoven et al. 2005). Genome sequencing has revealed a subset of arthropod genes encoding enzymes with clear selective advantages for herbivory. Studies on host adaptation in herbivorous arthropods have primarily focused on detoxification-related enzymes, such as cytochrome P450 monooxygenases, carboxyl/cholinesterases (CCEs), glutathione *S*-transferases, UDP-glycosyltransferases (UGTs), and ATP-binding cassette (ABC) transporters (Després et al., 2007; Dermauw and Van Leeuwen, 2014; Heidel-Fischer and Vogel, 2015). These families are widely conserved and have evolved primarily through vertical gene transfer and subsequent gene duplication. However, some arthropod detoxification genes show high similarity to microbial genes and unusual phylogenetic patterns, suggesting acquisition through HGT (Wybouw et al., 2016). Spider mites can thrive on plants containing cyanogenic glucosides, a major defense compound, as they express the *β-cyanoalanine synthase* acquired through HGT from bacteria (Wybouw et al., 2012, 2014, 2018). *DOG*s examined in the present study are also acquired through HGT from fungi (Njiru et al. 2022). The TSSM genome contains 17 *DOG* genes (Grbić et al., 2011; Dermauw et al., 2013). Some *DOG*s are expressed in the midgut and are induced in response to plant jasmonic acid-induced defense signaling (Njiru et al., 2022). Upon host plant change to tomatoes and in multi-pesticide resistant strains of TSSM, more than half of the *DOG* family members were differentially expressed, showing highly correlated expression patterns between tomato-adapted mites and pesticide-resistant strains (Dermauw et al., 2013). RNAi-mediated silencing confirmed the functional importance of *DOG*s in host plant utilization and pesticide resistance in TSSM (Xu et al., 2019; Njiru et al., 2022).

Biochemical characterization revealed differences in substrate specificity among some DOGs (Njiru et al., 2022). High cleavage activities were observed in TuDOG1 toward ellagic acid, TuDOG15 toward procyanidins A1 and B2, and TuDOG16 toward protocatechuic acid and caffeoyl malic acid. While Njiru et al. (2022) reported high cleavage activity of TuDOG15 toward EC, our study demonstrated that TkDOG15 and TuDOG15 also exhibited specific activity toward EGCg, EGC, and ECg in addition to EC (Supplemental Table 4-1). Furthermore, TkDOG15 showed significantly higher activity than TuDOG15 toward EGCg, EGC, and EC (Supplemental Table 4-1). When normalized by protein abundance in mites fed on tea (Figure 3e), TkDOG15 demonstrated distinctly higher activity toward all four catechins, and this difference was most pronounced for EC (Figure 4c). Consistently, tea-adapted KSM showed markedly lower sensitivity to EC in the toxicity assay, with the LC_50_ value exceeding the highest concentration tested (Figure 2b, Supplemental Table 2-1). The interspecific difference in DOG15 activity toward green tea catechins can be attributed both to differences in protein abundance and two amino acid substitutions (Q127A and T203A; Figure 4a, b). T203A is adjacent to the active-site residue Y202 and may contribute more directly to catalytic efficiency. Whether the two substitutions act individually or jointly remains to be determined, and dissecting their contributions by site-directed mutagenesis is a direction for future work. Interestingly, the amino acid sequence of TkDOG15 was identical between tea-adapted KSM and non-adapted KSM, suggesting that intraspecific differences in the contribution of DOG15 to tea adaptation mechanisms may be solely attributed to transcriptional regulation. RNAi-mediated silencing of the *TkDOG15* gene reduced the survival of tea-adapted KSM on tea leaves (Figure 3f), confirming the importance of enzyme abundance to tea adaptation. However, the reduction in survival was less pronounced than that observed in non-adapted KSM on tea leaves (Figure 1c). This discrepancy is likely due to ∼25% residual expression of the *TkDOG15* gene upon an RNAi treatment (Figure 3g). In addition, the delivery of dsRNA to tea-adapted KSM was limited to a 2-h oral administration in this study, leading to transient RNAi gene silencing. Further reduction in mite survival would be expected with sustained suppression of *TkDOG15* expression. Moreover, other detoxification-related enzymes were found to be upregulated in the transcriptome and proteome of tea-adapted KSM when fed on tea leaves (Supplemental Figure 3-1). Transcriptome analysis revealed elevated expression of three CCEs, two UGTs, and one ABC transporter (Supplemental Table 3-1). Proteome analysis identified increased abundance of one CCE and one UGT, as well as DOG1, DOG4, and DOG7 (Supplemental Table 3-2). Whether and how these detoxification enzymes contribute to the detoxification of green tea catechins or other tea defensive metabolites remains to be tested in future studies. Several other questions also remain open. First, the molecular basis of reduced chemosensory responses to catechins in tea-adapted KSM is unknown. Second, the regulatory mechanisms controlling DOG15 upregulation require further investigation. Third, the metabolic products of DOG15-mediated catechin degradation need characterization. Confirming that these products are less toxic than the parent catechins would further support the proposed detoxification role of DOG15. Finally, whereas DOG15 was the only enzyme upregulated at both the mRNA and protein levels in tea-adapted KSM on tea, the other enzymes, including DOG1, DOG4, and DOG7, as well as CCEs, UGTs, and ABC transporters, were enriched in either the transcriptome or the proteome. This suggests that tea adaptation in KSM may be multigenic, with these enzymes potentially acting synergistically with DOG15. Their contributions to catechin detoxification warrant functional validation.

In conclusion, we have identified key molecular mechanisms underlying adaptation to green tea in spider mites. Two amino acid substitutions in the horizontally transferred DOG15 enzyme, combined with upregulated expression, enable tea-adapted populations of Kanzawa spider mites to efficiently detoxify catechins and overcome their toxicity. In addition, tea-adapted KSM exhibit reduced contact irritancy and antifeedancy to catechins, though the molecular basis of these chemosensory changes remains unknown. These adaptations distinguish tea-adapted KSM from non-adapted populations and the super generalist TSSM. Our findings demonstrate that horizontal gene transfer followed by structural refinement and transcriptional upregulation can enable herbivores to rapidly exploit chemically defended plants. Over longer evolutionary timescales, such host plant-specific adaptations could contribute to reproductive isolation between populations and potentially drive ecological speciation in spider mites. These results advance our understanding of plant-herbivore coevolution and have practical implications for integrated pest management in tea cultivation.

## Materials and Methods

### Mites

A tea-adapted field population of the Kanzawa spider mite (KSM), *Tetranychus kanzawai* Kishida (Trombidiformes: Tetranychidae), was collected in a tea farm (Ureshino, Saga, Japan). A non-adapted KSM field population was collected in an apple orchard (Sobetsu, Hokkaido, Japan). A laboratory population of two-spotted spider mite (TSSM), *T. urticae* Koch, was obtained from the Western University (London, Ontario, Canada) (Grbić et al., 2011). Tea-adapted KSM was reared on leaves of kidney bean (*Phaseolus vulgaris* L.) and tea (*Camellia sinensis* (L.) Kuntze). Non-adapted KSM and TSSM were reared on kidney bean leaves.

### Mite performance test

Tea-adapted KSM, non-adapted KSM, and TSSM were grown on kidney bean leaves until the teleiochrysalid stage. Fifteen newly-emerged adult females from each population were transferred to a single tea leaf and reared at 25 °C for 10 days. For the performance test using kidney bean, newly-emerged adult females of tea-adapted KSM and TSSM were transferred to a single bean leaf. The number of survivors and eggs laid on each leaf were assessed daily. Mites that escaped from leaves during the experiment were excluded from the data. Data were collected from 5 independent experimental runs (*n* = 15 per run).

### HPLC analysis

We used an HPLC system consisting of a PU-980 pump (JASCO Corp., Tokyo, Japan), UV-970 Intelligent UV VIS Detector (JASCO), and a J-Pak Wrapsil C18 column (JASCO; 250 mm × 4.6 mm, 5 µm). A tea leaf disc (10 mm in diameter) was placed in a 1.5 mL tube and homogenized under liquid nitrogen using a homogenizer pestle. An extraction buffer of 50% (v/v) acetonitrile (ACN) in water was added to the tube, and the green tea catechins were extracted from the sample for 1 h at room temperature using a tube tumbler rotator (SBS-550; Labnet International Inc., Edison, NJ, USA). After extraction, the solid fraction removed by centrifugation at 21,500 ×g at 25 °C for 5 min using a spin column (Ultrafree-MC Centrifugal Filter UFC30LG00; Merck KGaA, Darmstadt, Germany), and the filtrate was analyzed by HPLC. The standard solutions of EGCg (CAS No. 989-51-5; FUJIFILM Wako Pure Chemical Corp., Osaka, Japan), EGC (CAS No. 970-74-1; FUJIFILM Wako Pure Chemical), ECg (CAS No. 1257-08-5; FUJIFILM Wako Pure Chemical), EC (CAS No. 490-46-0; FUJIFILM Wako Pure Chemical), and caffeine (CAS No. 58-08-2; FUJIFILM Wako Pure Chemical) were dissolved in the buffer to make a standard solution, respectively. Each 5 µL of sample or standard solution was injected into the HPLC system using a glass micro-syringe (GL Sciences Inc., Tokyo, Japan). The analytical conditions were a flow rate of 1 mL/min and a detection wavelength of 280 nm. A gradient elution was performed using solvent A (Water: ACN: HPO_4_ = 499.5: 500: 0.5) and solvent B (Water: ACN: HPO_4_ = 954.5: 45: 0.5) as the mobile phase. Data were collected from 5 independent experimental runs.

### Toxicity assay

A piece of Kimwipe (S-200; Nippon Paper Crecia Co., Ltd., Tokyo, Japan) was infiltrated with an aqueous solution containing individual catechins (EGCg, EGC, ECg, and EC) or a mixture of the four catechins (NH025104; Nagara Science Co., Gifu, Japan) at concentrations of 0 (control), 10, 10^2^, 10^3^, 10^4^, and 10^5^ ppm (w/v). This range was chosen to encompass the individual catechin levels in fresh tea leaves (Figure 2a). The solution was then mixed with 1% (w/v) brilliant blue FCF (FUJIFILM Wako Pure Chemical). According to Bensoussan et al. (2020), newly-molted adult females of tea-adapted KSM, non-adapted KSM, and TSSM were incubated on the infiltrated Kimwipe piece at 25 °C for 24 h. After 24 h, adult females whose midguts had turned blue due to pigmentation were selected under a stereomicroscope. This pigmentation was visible due to the semi-transparent epidermis (Ghazy et al., 2020). Adult females were then transferred to a polystyrene Petri dish with a diameter of 55 mm (AGC Techno Glass Co., Ltd., Shizuoka, Japan). Water-soaked Kimwipes were placed on the inner sides of the dish to prevent mite escape. The number of survivors was counted after 3 days. Data were collected from 3 to 5 independent experimental runs (*n* = 25 to 45 per run).

### Irritancy assay

One-half (4.5 × 9.0 mm) of the area on one side of a 9.0 × 9.0 mm glass plate (Matsunami Glass Ind., Ltd., Osaka, Japan) was coated with the catechin mixture at a concentration of 10^5^ ppm (=10%) (w/v) or 10 mM fenpropathrin (CAS No. 39515-41-8; FUJIFILM Wako Pure Chemical) in a solvent consisting of 0.1% (w/v) brilliant blue FCF and 0.1% (v/v) Tween 20 in methanol. Fenpropathrin was used as a positive control due to its known TSSM-proofing effect (Penman et al., 1986). The other half of the plate was coated with the solvent. For the negative control, the entire area on one side of the plate was coated with the solvent. Newly-molted adult females of tea-adapted KSM, non-adapted KSM, and TSSM were placed on the glass plates. The locomotion of each test mite on the plate was recorded for 10 min at 29.97 fps using a digital camera (EOS Kiss X7; Canon Inc., Tokyo, Japan) installed on a stereomicroscope (M205FA or S8 APO; Leica Microsystems GmbH, Wetzlar, Germany). We then used a Python program (Hamdi et al., 2023) to calculate the number of frames for which a mite was located in the treatment region coated with the catechin mixture or fenpropathrin and in the solvent region. Data from mites with an average locomotion speed of less than 0.30 mm/s over 10 min or those that escaped from the glass plate were excluded from the analyses, because inactive mites showed insufficient movement to determine chemo-orientation behavior and escaped mites did not complete the assay. To quantify chemo-orientation behavior, we used the following equation (Sakuma and Fukami, 1985): *PI* = (*N*_T_ − *N*_S_)/(*N*_T_ + *N*_S_), where *PI*, *N*_T_, and *N*_S_ indicate the preference index, the number of frames of the mite located in the treatment region, and the number of frames of the mite located in the solvent region, respectively. The *PI* value ranges from −1 (complete localization in the solvent region) to +1 (complete localization in the treatment region). Different individuals (*n* = 10 to 11) were used for each round.

### Antifeedancy assay

Adaxial surface of leaf discs (10 mm in diameter) of kidney bean were coated with solutions of the catechin mixture at a concentration of 10^5^ ppm (w/v) or 10 mM fenpropathrin (positive control) in methanol. For coating, 7 µL of the test solution or solvent (negative control) was dropped onto a piece of nylon mesh sheet (100 µm in opening; As One Corp., Osaka, Japan) covering the adaxial surface of each leaf disc. The droplet spread across the entire surface of leaf disc due to the surface tension of the mesh sheet, even without surfactants. After 1 min of coating, the mesh was removed. Leaf discs were then covered with stretched Parafilm (Parafilm M; Bemis Co. Inc., Neenah, WI) that prevented direct contact of mites’ legs with test compounds distributed on the surface of leaf discs but allowed mites to penetrate with their stylets for feeding on the leaf discs. To prevent mite from escaping, water-soaked Kimwipes were placed on the stretched Parafilm except the arena directly above the test leaf disc, which was accessible for mites to feed. We used this setup for evaluating the antifeedant effect of compounds on mites without irritancy caused by chemosensing by legs. Each 15 newly-molted adult females of tea-adapted KSM, non-adapted KSM, and TSSM were placed in the feeding arena of each setup. After 24 h, all females were removed from the setup. The covered Parafilm was removed from each leaf disc and a RGB image of the adaxial surface was acquired using an image scanner (GT-X980; Seiko Epson Corp., Nagano, Japan). According to Cazaux et al. (2014), damaged area by mite feeding was analyzed from the RGB image of each leaf disc using Adobe Photoshop CC 2023 (Adobe Inc., San Jose, CA). Data were collected from 3 to 7 independent experimental runs (*n* = 15 per run).

### Transcriptome analysis

Newly-molted adult females of tea-adapted KSM, non-adapted KSM, and TSSM were transferred onto leaves of tea plants and kidney bean seedlings. To limit the dispersal of mites to other leaves, water-soaked Kimwipe pieces were wrapped around each petiole of leaves on which mites were transferred. After 24 h, each 150 mites were collected. The samples frozen in liquid nitrogen were then crushed using a homogenizer pestle. Total RNA was extracted using a NucleoSpin RNA Plus XS kit (Macherey-Nagel GbmH & Co. KG, Düren, Germany) according to the manufacturer’s protocol. RNA purity and concentration were measured using a spectrophotometer (NanoPhotometer N60; Implen GmbH, Munich, Germany). The cDNA libraries were prepared with the TruSeq Stranded mRNA (Illumina Inc., San Diego, CA, USA). The transcriptomes were analyzed in paired ends (2 × 150 bp) using a NovaSeq 6000 system (Illumina). Raw sequence data were generated by the Illumina pipeline and fastp (v.0.23.2; Chen et al., 2018) was used to filter out Illumina adapter, reads containing more than 5% unknown nucleotides and low-quality reads (reads with Q-values less than 20) to clean reads were generated. Using the transcriptome data of TSSM (Grbic et al., 2011) as a reference, clean reads quantified tea-adapted KSM, non-adapted KSM, and TSSM on tea and kidney bean leaves. Kallisto (ver. 0.46.2; Bray et al., 2016) was used to quantify clean reads, and count and transcripts per million (TPM) data were obtained for each gene. The bootstrap count was set to 100. Relative Log Expression (RLE) normalization with DESeq2 (Love et al., 2014) via iDEP (ver. 0.93; Ge et al., 2018) was performed from the count data. Genes with an adjusted *p*-value < 0.05 and a log2 fold change ≥ 1 were selected as differentially expressed genes. These criteria follow those commonly applied in mite transcriptomic studies (Vidal-Quist et al., 2025). Data were collected from two independent experimental runs (*n* = 150 per run).

### Proteome analysis

Mite samples were collected using the same protocol as for the transcriptome analysis. The samples frozen in liquid nitrogen were crushed using a homogenizer pestle and stored at −80 °C until protein extraction. Protein lysis buffer consisting of 10 mM Tris-HCl (pH 9.0) and 8 M urea was added to tube containing each sample. The samples were further crushed using a homogenizer pestle and a sonicator (AS38A; As One). Each tube was centrifuged at 13,760 ×g at 4 °C for 10 min to remove debris, and the supernatant was used as the following samples. The protein concentration was measured using a Qubit 4 fluorometer (Thermo Fisher Scientific Inc., Waltham, MA, USA). Each sample containing 40 μg of protein was reduced with 10 mM DTT (Invitrogen) for 30 min and then alkylated with 50 mM IAM (CAS No. 144-48-9; Fujifilm Wako Pure Chemical) for 20 min under shade. Then each sample was diluted with 50 mM NH_4_HCO_3_ (CAS No. 1066-33-7; Fujifilm Wako Pure Chemical) to a quarter urea concentration, followed by digestion into peptides overnight with trypsin (Promega K.K., Tokyo, Japan) at a ratio of 1:50 (trypsin: protein) at room temperature. To stop the trypsin reaction, 2% TFA was added to the sample at the same volume and the solution was centrifuged at 13,000 rpm and 4 °C for 10 min. Solution A (water: ACN: TFA = 200: 800: 1) was added to a 200 µL of peptide desalting column (GL-Tip SDB; GL Sciences) and centrifuged at 1,100 ×g and 4 °C for 3 min. Next, solution B (water: ACN: TFA = 950: 50: 1) was added under the same conditions. The sample was then dropped into the column and washed with solution B. Then, the sample was eluted with solution A. The samples were concentrated in an evaporator under vacuum and stored at −80 °C until they were applied to an LC-MS/MS (Orbitrap Exploris 480; Thermo Fisher Scientific). The peptides were identified using Proteome Discoverer v.2.5.0.400 (Thermo Fisher Scientific) with raw data. The protein database was TSSM annotation (v.20190125) from the ORCAE database (Sterck et al., 2012; https://bioinformatics.psb.ugent.be/orcae/overview/Tetur). The relevant parameters were as follows: “trypsin” for the enzyme name; “2” for the maximum number of missed cleavage sites; “20 ppm” for mass tolerance; “0.05 Da” for fragment mass tolerance; and “1%” for the false discovery rate. Differentially expressed proteins were detected based on a *p*-value < 0.05 and a log2 fold change ≥ 1. These criteria are consistent with those used in our previous spider mite proteomic analysis (Arai et al., 2025). Data were collected from three independent experimental runs (*n* = 100 to 150 per run).

### dsRNA synthesis

Total RNA was extracted from about 300 adult females of tea-adapted KSM. Then, cDNA was synthesized from 5 µg of total RNA using reverse transcriptase (SuperScript IV Reverse Transcriptase; Thermo Fisher Scientific), 50 µM oligo d(T)_20_ primer (Thermo Fisher Scientific), and 10 mM dNTP mix (10 mM each) according to the manufacturer’s protocol, and stored at −30 °C. Genomic DNA (gDNA) was extracted from about 300 adult females of tea-adapted KSM using a Quick-DNA Microprep kit (ZYMO Research, Irvine, CA, USA) according to the manufacturer’s protocol and stored at 4 °C. Using cDNA or gDNA as a template, specific primers targeting a 568-bp fragment of the *TkDOG15* gene (ortholog of *tetur20g01790*) or a 382-bp fragment of the intergenic region (negative control (NC), genomic coordinates: scaffold 12, position 1690614–1690995; Suzuki et al. 2017a) were used, respectively, for PCR amplification using KOD-Plus DNA polymerase (Toyobo Co. Ltd., Osaka, Japan). Primers designed to amplify the DNA fragments of *TkDOG15* and NC are shown in Table 1. PCR fragments were purified with NucleoSpin Gel and PCR Clean-Up Kit (Macherey-Nagel). A template of 100 ng of each PCR fragment was used for RNA synthesis using T7 RNA Polymerase (*in vitro* Transcription T7 Kit; Takara Bio Inc., Kusatsu, Japan) according to the manufacturer’s protocol. Then DNase I (Takara Bio) treatment for 30 min at 37 °C, RNA was denatured at 95 °C, followed by slow cool-down to room temperature to facilitate dsRNA formation. The dsRNA fragments (ds*TkDOG15* and dsNC) were purified using phenol-chloroform and precipitated with 99.5% ethanol. The concentration and size of dsRNA was measured with the spectrophotometer and 1.5% (w/v) and agarose gel electrophoresis (Agarose 21; Nippon Gene Co. Ltd., Tokyo, Japan), respectively.

**Table 1.**
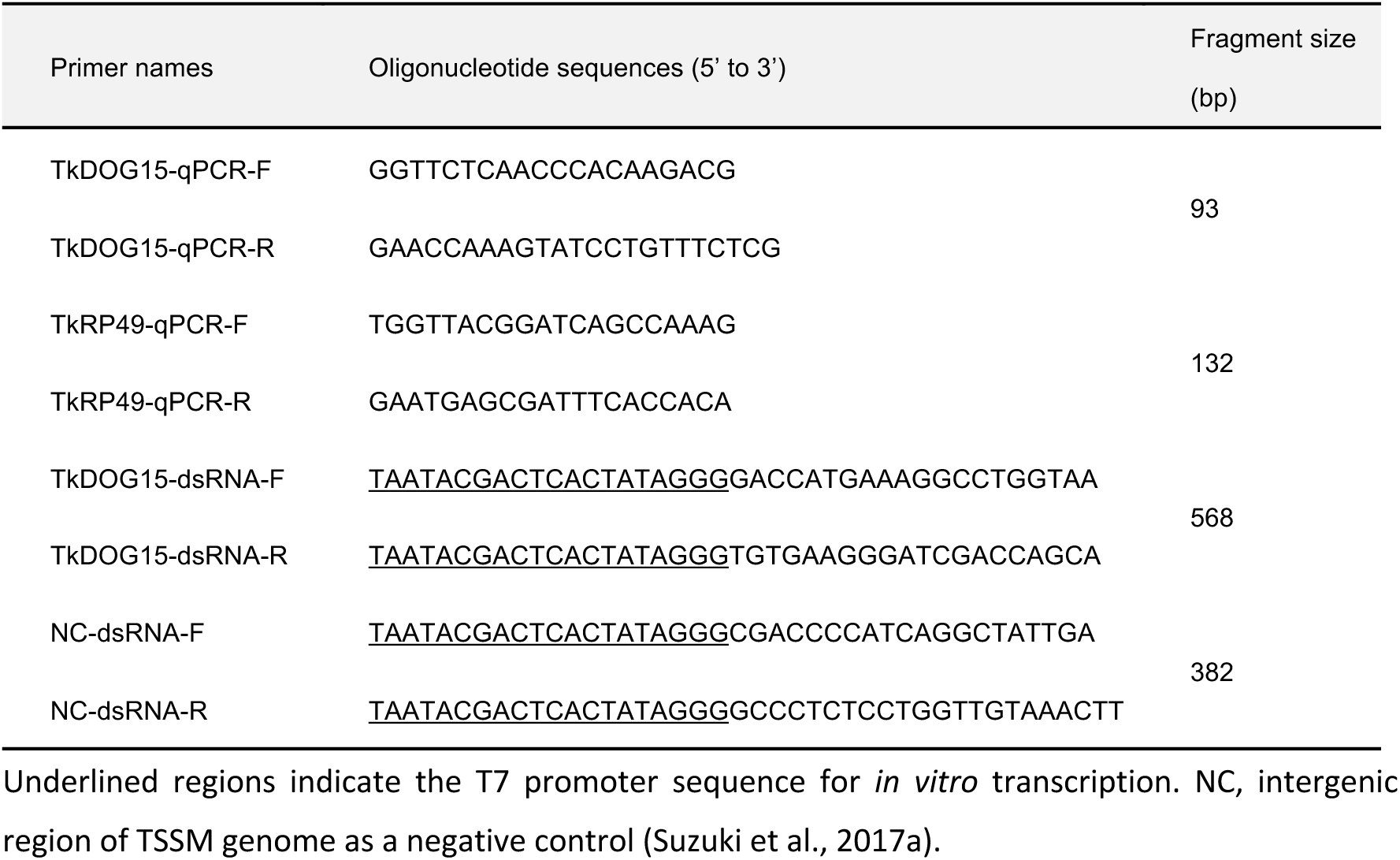
Primers for real time qRT-PCR and dsRNA synthesis.

### RNAi

According to Suzuki et al. (2017a, b) and Bensoussan et al. (2020), newly-molted adult females of tea-adapted KSM were transferred to a 1 cm^2^ piece of Kimwipe infiltrated with ds*TkDOG15* or dsNC at 1 µg/µL in 1% (w/v) brilliant blue FCF (FUJIFILM Wako Pure Chemical). Mites were incubated at 25 °C for 2 h to orally deliver dsRNA. Mites with blue pigmentation in their midgut were selected under a stereomicroscope and transferred onto tea plants (15 mites/leaf). The number of survivors and eggs laid was assessed every day. Data were collected from five independent experimental runs (*n* = 15 per run).

### Real-time quantitative reverse transcription-PCR (qRT-PCR) analysis

Adult females of tea-adapted KSM were collected at 4 days after feeding on ds*TkDOG15* or dsNC, kept frozen in liquid nitrogen, and stored at −80 °C. Total RNA was extracted from 50 adult females in each experiment, and single-stranded cDNA was synthesized by reverse transcription of total RNA using a High Capacity cDNA Reverse Transcription Kit (Applied Biosystems). qRT-PCR reactions were performed in three independent experimental runs (*n* = 40 to 50 mites per run) with Power SYBR Green Master Mix (Applied Biosystems) on an ABI StepOnePlus Real-Time PCR System (Applied Biosystems). A gene encoding a ribosomal protein, *RP49*, was used as a reference gene (Suzuki et al., 2017a). Primers for the reference gene and target gene are shown in Table 1. The threshold cycle (*Ct*) value was calculated by averaging three technical replicates. The expression value of the target gene was normalized to the reference gene. The normalized relative quantity (*NRQ*) was calculated as follows: *NRQ* = (1 + *E*_R_)*^Ct^*^R^/(1 + *E*_T_)*^Ct^*^T^, where E_R_ and E_T_ are the amplification efficiencies for reference and target genes, respectively. *Ct*R and *Ct*T are the cycle threshold values of reference and target genes, respectively.

### Protein production, purification and activity assay

The sequence for *TuDOG15* (*tetur20g01790*) was obtained from the ORCAE database. The sequence for *TkDOG15* was obtained through RNA-seq reads. The predicted signal peptide at the N-terminal was not included when generating constructs. The constructs containing the mature sequence of *TuDOG15* and *TkDOG15* were ordered from SynBio (Monmouth Junction, NJ) in pET30a(+) vectors. Plasmids were transformed into *E. coli* (DH5α) through the heat shock method, then spread on 2% agar plates containing 50 µg/mL of kanamycin. The plasmid for each construct was purified from single colonies with the GeneJET Plasmid Miniprep Kit (Thermo Fisher Scientific, Waltham, MA) and used to transform *E. coli* (BL21(DE3)) cells using the previous method. A single colony was picked and grown in 5 mL LB medium overnight with 50 µg/mL of kanamycin and 50 mg of glucose. After 16 hours, the 5 mL cultures were transferred into 1L of Luria Broth (LB) medium with the same kanamycin concentration as used prior. The flasks containing the 1L cultures were then incubated at 37 °C and 250 rpm until the OD_600_ reached 0.8. The cultures were cooled to 16 °C and induced with 0.4 mM isopropyl-β-D-thiogalactopyranoside (IPTG) for protein expression. During expression, 3 mg of ferrous sulphate was added to increase the incorporation of the iron cofactor for the overexpressed protein. The cultures continued protein production overnight at 16 °C, shaking at 150 rpm. Cells were centrifuged down for 15 minutes at 4 °C and 13,000 g. The supernatant was discarded for all cultures, and the pellets were collected and stored at −80 °C until used.

Both TkDOG15 and TuDOG15 were purified using the same protocol. Cell pellets were thawed and resuspended with lysis buffer (50 mM Tris, 500 mM NaCl, 10 mM imidazole, 20 mM β-mercaptoethanol, 2% glycerol (v/v), pH 7.5) at a ratio of 5 mL buffer per 1 gram of pellet. The resuspended solution was lysed via sonication for 10 cycles, with each cycle having 10 seconds sonication and 50 seconds rest at 250 W. The cell debris was removed through centrifugation for 40 minutes at 4 °C and 36,000 g. The supernatant was collected and used for purification. First, ammonium sulfate ((NH_4_)_2_SO_4_) precipitation was used. The supernatant was stirred slowly for 45 minutes at 4 °C with ((NH_4_)_2_SO_4_) steadily added until the solution reached 25% ammonium sulfate. Following, the solution was centrifuged down for 30 minutes at 4 °C and 3082 g. The mixture was then stirred again with the same parameters but changing the final saturation to 50% ammonium sulfate. This was followed by centrifugation as previously mentioned. The pellet remaining from centrifuging was resuspended using phosphate buffer saline and dialyzed overnight in 50 mM Tris, pH 7.5 at 4 °C. After 16 hours, the sample was injected on a HiPrep DEAE FF 16/1 anion exchange column (Cytiva, Marlborough, MA). The column was equilibrated with 50 mM Tris, pH 7.5, and eluted using a 0-40% linear gradient with 50 mM Tris, 2 M NaCl, pH 7.5. The protein was then collected and injected onto a HiLoad 16/600 Superdex 200 pg size exclusion column (Cytiva, Marlborough, MA). The protein content and sample purity were verified on a 12% SDS-PAGE. The protein concentration was determined on a NanoDrop 2000c Spectrophotometer (Thermo Fisher Scientific, Waltham, MA) using the molecular weight and extinction coefficient predicted by ExPASy ProtParam (Wilkins, et al. 1999). A ferrozine assay was performed to test the iron concentration for TkDOG15 and TuDOG15 using the previously described protocol (Schlachter, et al. 2019).

To test the activity of TkDOG15 and TuDOG15, the four catechins tested (EC, ECg, EGC, and EGCg) were purchased from Cayman Chemicals (Ann Arbor, MI). A Hansatech Clark-type oxygen electrode (Oxytherm, Hansatech) was used to measure the oxygen consumed from the enzyme catalyzed reaction. The specific activity was tested with 125 µM of each catechin, mixed with either 1.5 µM (for EC and ECg) or 4.5 µM (for EGC and EGCg) of TkDOG15 or TuDOG15. This reaction was carried out in 500 µL of 25 mM bis-Tris, 150 mM NaCl, pH 6.0. The solution maintained 25 °C and stirred at 70 rpm by the Oxytherm chamber. The rate of oxygen consumption was measured for 20 seconds. The specific activities were corrected for active, iron-bound enzyme for each set of reactions.

### Statistical analysis

All data were analyzed using R v.4.2.2. Survivorship in adult females of spider mites on tea and kidney bean, and in tea-adapted KSM on tea after RNAi-mediated silencing of the *TkDOG15* gene were plotted with the Kaplan–Meier method (*survfit*() function in the R package ‘*survival*’). Differences in curves among three populations on tea, between tea-adapted KSM and TSSM on tea and kidney bean, and between ds*TkDOG15* and dsNC in tea-adapted KSM were analyzed by the log-rank test (*survdiff*() function in the R package ‘*survival*’). Dose–response relationships between catechin levels and survivorship of spider mites were determined, and curves were generated with the two-parameter log-logistic function (*drm*() function in the *R* package ‘*drc*’); they are presented with a 95% confidence interval. Fecundity of adult females of three populations on tea, and mRNA and protein expression levels of DOG15 in three populations on tea and kidney bean leaves were arcsine-square-root transformed, and the normalized data were analyzed by ANOVA (*aov*() function in the R package ‘*stats*’) followed by Tukey’s HSD test (*glht*() function in the R package ‘*multcomp*’). Fecundity of adult females of tea-adapted KSM and TSSM on tea and kidney bean, relative expression level of the *TkDOG15* gene in tea-adapted KSM after the RNAi treatment with ds*TkDOG15* and dsNC, and relative enzymatic activity between TkDOG15 and TuDOG15 were analyzed with Student’s *t*-test (*t.test*() function in the R package *‘stats’*). The preference index in three populations by chemo-orientation behavioral assay and feeding preference in three populations fed kidney bean coated treatments were analyzed with Dunnett’s test (*glht*() function in the R package ‘*multcomp*’).

## Acknowledgments

The authors would like to thank Mr. N. Hayashi (OAT Agrio Co., Ltd., Tokyo, Japan), Dr. Y. Sato (Institute of Fruit Tree and Tea Science, NARO, Japan), Dr. T. Gotoh (Ibaraki University, Ibaraki, Japan), and Dr. M. Grbić (The University of Western Ontario, London, ON, Canada) for providing the mite populations used in this study. This work was supported by JSPS KAKENHI grant 22J22156 awarded to NT. This work was also supported in part by JSPS KAKENHI grants 16K18661, 18H02203, 21H02193, and 24K21256, as well as the Cabinet Office, Government of Japan, Cross-ministerial Moonshot Agriculture, Forestry and Fisheries Research and Development Program, “Technologies for Smart Bio-industry and Agriculture” (JPJ009237; funding agency: Bio-oriented Technology Research Advancement Institution, Kawasaki, Japan), awarded to TS. This work was also supported by USDA’s National Institute of Food and Agriculture (award # 2020–67014-31179 for MC and VG), through the NSF/NIFA Plant Biotic Interactions Program.

## Author contributions

Conceptualization, NT, BA, VZ, VG, MC, and TS; methodology, NT, BA, RHA, RM, SS, MY, MC, and TS; investigation, NT, BA, RHA, and RM; formal analysis, NT, BA, and RHA; visualization, NT, BA, and RHA; writing – original draft, NT and TS; writing – review and editing, NT, BA, RHA, RM, SS, MY, VZ, VG, MC, and TS; supervision, VG, MC, and TS; funding acquisition, NT, MC, and TS.

## Data Availability Statements

RNA-seq data have been deposited at BioProject (accession: PRJNA1356475). Proteomics data have been deposited at jPOSTrepo (accession: JPST004156) and ProteomeXchange (accession: PXD070535). These data are publicly available as of the date of publication.

## Additional files

**Supplemental Figure 1-1**

The caption is included within the file.

**Supplemental Figure 1-2**

The caption is included within the file.

**Supplemental Figure 3-1**

The caption is included within the file.

**Supplemental Figure 4-1**

The caption is included within the file.

**Supplemental Table 2-1**

The caption is included within the file.

**Supplemental Table 3-1**

The caption is included within the file.

**Supplemental Table 3-2**

The caption is included within the file.

**Supplemental Table 4-1**

The caption is included within the file.

**Supplementary Movie 1**

Locomotor behavior of tea-adapted KSM on the control treatment (10 min, 32×).

**Supplementary Movie 2**

Locomotor behavior of tea-adapted KSM exposed to the catechin mixture (10 min, 32×).

**Supplementary Movie 3**

Locomotor behavior of tea-adapted KSM exposed to fenpropathrin (10 min, 32×).

**Supplementary Movie 4**

Locomotor behavior of non-adapted KSM on the control treatment (10 min, 32×).

**Supplementary Movie 5**

Locomotor behavior of non-adapted KSM exposed to the catechin mixture (10 min, 32×).

**Supplementary Movie 6**

Locomotor behavior of non-adapted KSM exposed to fenpropathrin (10 min, 32×).

**Supplementary Movie 7**

Locomotor behavior of TSSM on the control treatment (10 min, 32×).

**Supplementary Movie 8**

Locomotor behavior of TSSM exposed to the catechin mixture (10 min, 32×).

**Supplementary Movie 9**

Locomotor behavior of TSSM exposed to fenpropathrin (10 min, 32×).

