## Supplemental Figure 1-1 for "Adaptation of an herbivorous arthropod to green tea plants by overcoming catechin defenses"

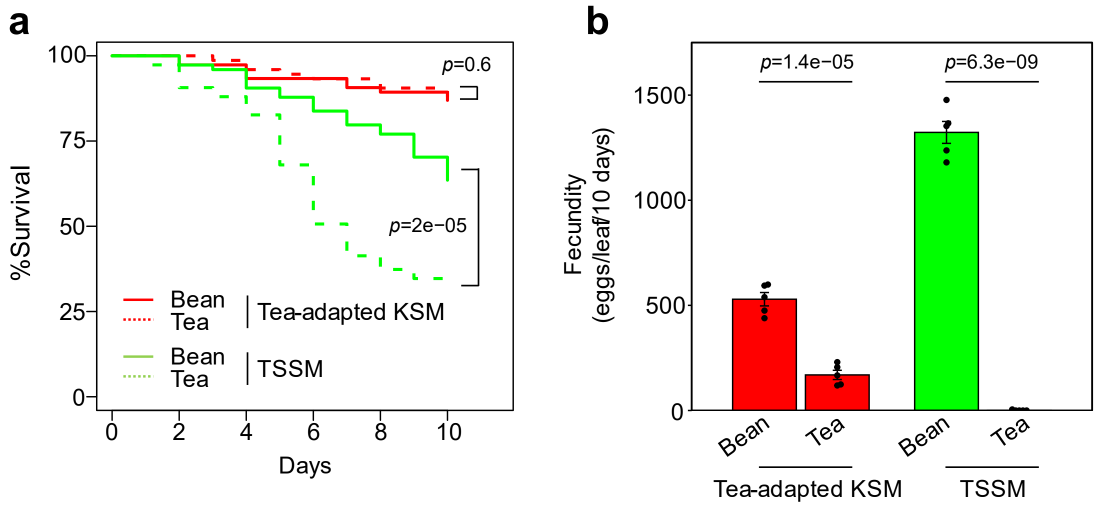


**Supplemental Figure 1-1. Mite performance on tea and bean leaves.**

(a) Survival and (b) fecundity in tea-adapted KSM and TSSM on tea and kidney bean leaves (15 mites/leaf) at 25 °C for 10 days. In each population, 15 mites that had grown to the teleiochrysalid stage on kidney bean leaves and molted to adults within 2 h were transferred to a single leaf of tea or kidney bean leaf and the numbers of survivors and eggs laid were counted daily (15 mites/leaf). Mites that escaped from the leaf during the experiment were excluded from the data; the total of 70 to 75 mites in each treatment were used for the survival analysis. Fecundity data were collected from 5 independent experimental runs. Mite survival and fecundity were compared between on kidney bean leaf and on tea leaf using the log rank test and Student’s *t*-test, respectively.
