## Supplemental Figure 1-2 for "Adaptation of an herbivorous arthropod to green tea plants by overcoming catechin defenses"

Fig. S1-2

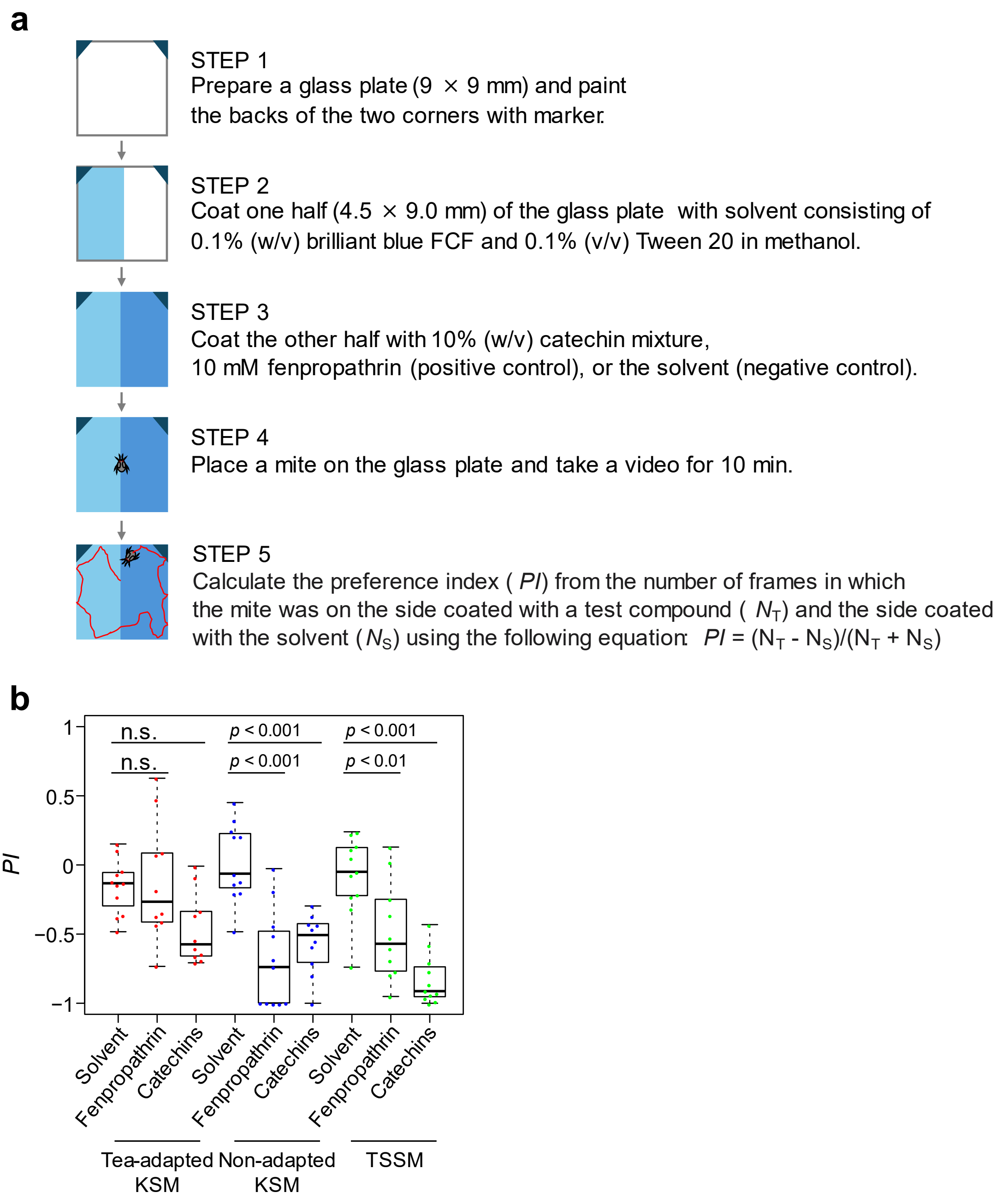


**Supplemental Figure 1-2. Description of the chemo-orientation behavioral assay procedure.**

(a) Schematic of the procedure for preparing the glass plate for the behavioral assay. Locomotion of tea-adapted KSM, non-adapted KSM, and TSSM on the glass plate (1 mite/plate) was recorded with a digital camera for 10 min. The location of the mite on the glass plate was analyzed from each frame of the video and a preference index (*PI*) was calculated. *PI* ranges from −1 (complete localization on the solvent coated side) to +1 (complete localization in the test compound coated side). (b) Chemo-orientation behavior of tea-adapted KSM, non-adapted KSM, and TSSM towards test compounds ($n$ = 10 to 11). *PI* was compared using Dunnett’s test.
