## Supplemental Figure 3-1 for "Adaptation of an herbivorous arthropod to green tea plants by overcoming catechin defenses"

Fig. S3-1

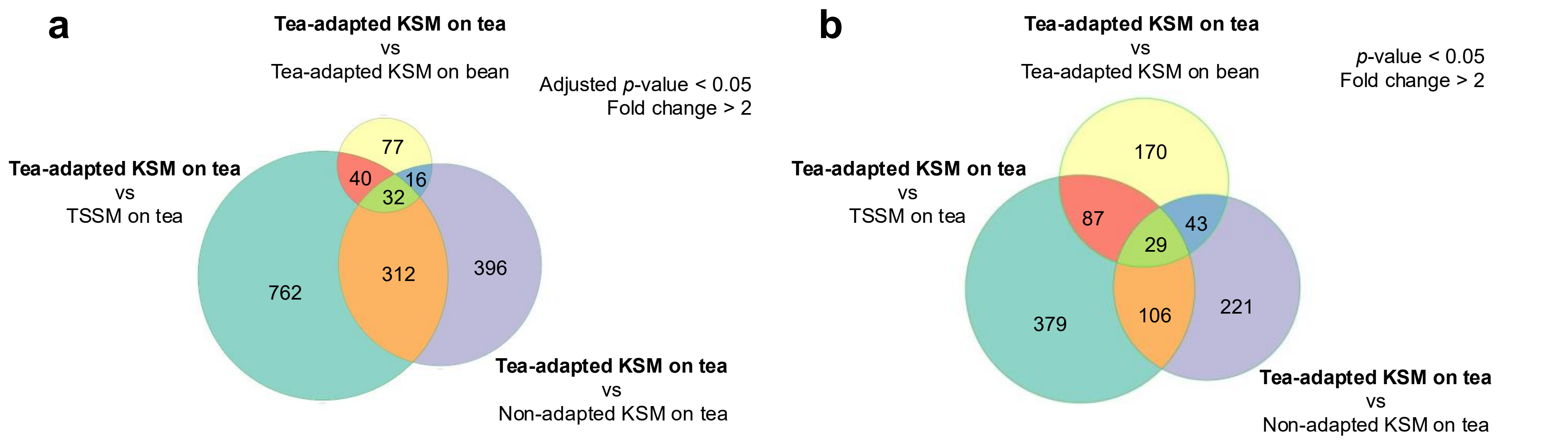


**Supplemental Figure 3-1. Number of highly expressed genes in (a) transcriptome and (b) proteome of tea-adapted KSM fed on tea leaves compared to tea-adapted KSM fed on bean leaves, non-adapted KSM fed on tea leaves, and TSSM fed on tea leaves.**

Comparative transcriptomics and proteomics were performed between tea-adapted KSM fed on tea leaves, tea-adapted KSM fed on kidney bean leaves, non-adapted KSM fed on tea leaves, and TSSM fed on tea leaves. Adult females were allowed to feed on each host plant at 25 °C for 24 h. Data were collected from 2 (transcriptome) and 3 (proteome) independent experimental runs with 100 to 150 adult females each. In tea-adapted KSM, 165 mRNAs and 329 proteins were highly expressed when fed on tea leaves compared to on bean leaves. When fed on tea leaves, 756 mRNAs and 399 proteins were highly expressed in tea-adapted KSM compared to non-adapted KSM. When fed on tea leaves, 1,146 mRNAs and 601 proteins were highly expressed in tea-adapted KSM compared to TSSM.
