## Supplemental Figure 4-1 for "Adaptation of an herbivorous arthropod to green tea plants by overcoming catechin defenses"

**Fig. S4-1**

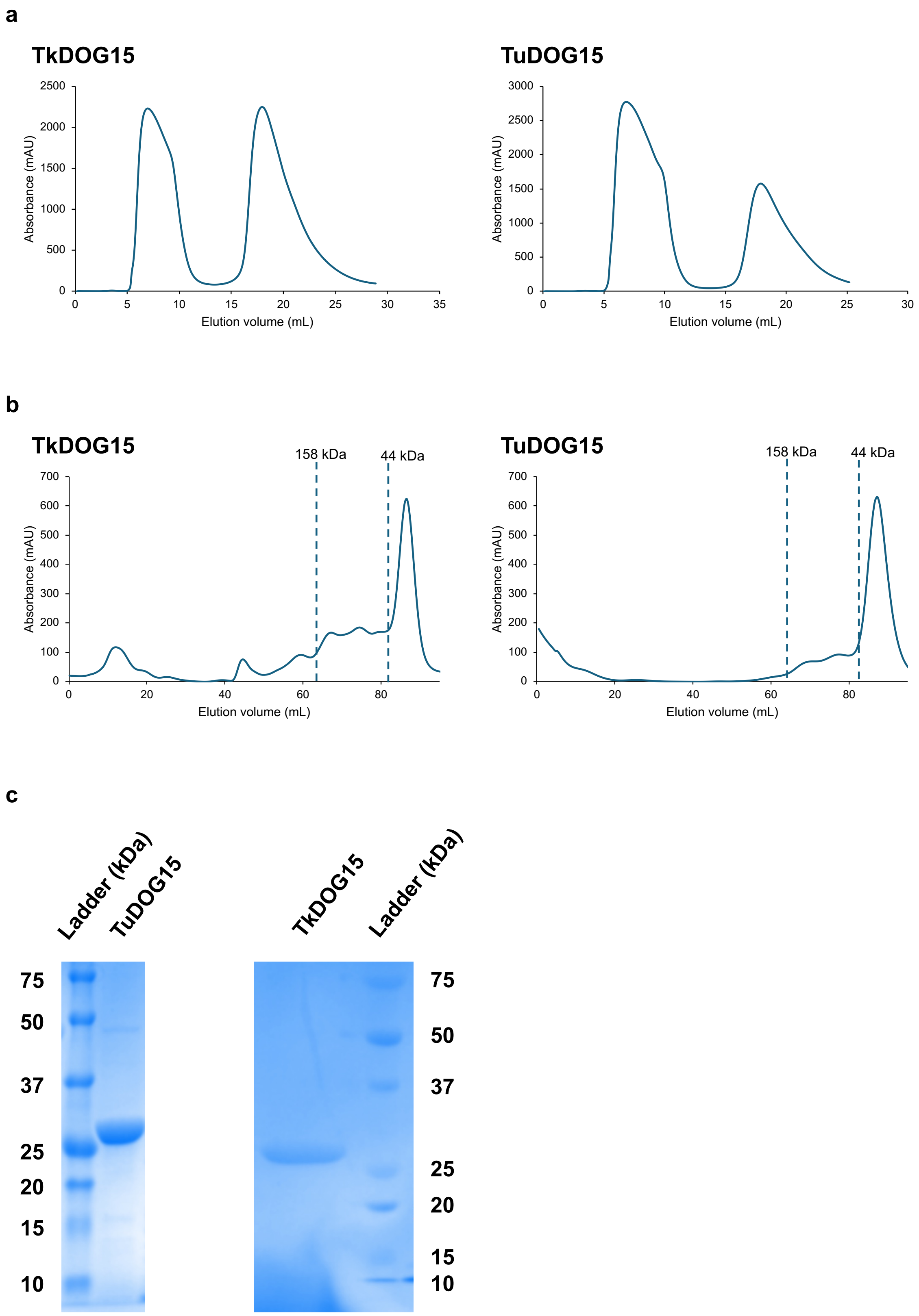


**Supplemental Figure 4-1. (a) Anion chromatograms, (b) size-exclusion chromatograms, and (c) SDS-PAGE gels for TkDOG15 and TuDOG15.**
