## Supplemental Table 2-1 for "Adaptation of an herbivorous arthropod to green tea plants by overcoming catechin defenses"

**Table S2-1**

**Supplemental Table 2-1. Lethal sensitivity to green tea catechins in tea-adapted KSM and TSSM.** Mites were soaked in different concentrations of an aqueous solution of (−)-epigallocatechin gallate (EGCg), (−)-epigallocatechin (EGC), (−)-epicatechin gallate (ECg), or (−)-epicatechin (EC) at 25 °C for 24 h.

| Catechins | Tea-adapted KSM | | TSSM | |
|---|---|---|---|---|
| | $LC_{50}$[a] (ppm) | 95% CI[b] (ppm) | $LC_{50}$ (ppm) | 95% CI[b] (ppm) |
| EGCg | $2.4 \times 10^3$ | $1.4 \times 10^3$ to $3.4 \times 10^3$ | $3.9 \times 10^3$ | $1.9 \times 10^3$ to $5.9 \times 10^3$ |
| EGC | $5.2 \times 10^4$ | $1.4 \times 10^4$ to $9.1 \times 10^4$ | $3.7 \times 10^3$ | $1.8 \times 10^3$ to $5.6 \times 10^3$ |
| ECg | $>10^5$[c] | – | $9.2 \times 10^2$ | $4.2 \times 10^2$ to $1.4 \times 10^3$ |
| EC | $>10^5$[c] | – | $1.2 \times 10^3$ | $6.6 \times 10^2$ to $1.7 \times 10^3$ |

[a]50% lethal concentration (lower to upper limits). [b]Confidence limits. [c]$LC_{50}$ values could not be determined within the tested concentration range (up to $10^5$ ppm).
