## Supplemental Table 3-1 for "Adaptation of an herbivorous arthropod to green tea plants by overcoming catechin defenses"

**Table S3-1**

**Supplemental Table 3-1. List of 32 mRNAs highly expressed in tea-adapted KSM fed on tea leaves.** The levels of mRNAs were compared to tea-adapted KSM fed on bean leaves, non-adapted KSM fed on tea leaves, and TSSM fed on tea leaves.

| Database ID | Gene annotation | Length (bp) | Log2 fold change | | |
|---|---|---|---|---|---|
| | | | vs Tea-adapted KSM on bean | vs TSSM on tea | vs Non-adapted KSM on tea |
| tetur01g10060 | G protein-coupled receptor, rhodopsin-like | 3201 | 1.03 | 2.68 | 1.72 |
| tetur01g10090 | Hypothetical protein | 951 | 1.38 | 12.23 | 7.08 |
| tetur01g14180 | Carboxyl/cholin esterase (CCE12) | 1683 | 3.81 | 2.53 | 1.00 |
| tetur01g91139 | Hypothetical protein | 636 | 5.52 | 3.67 | 5.52 |
| tetur02g92473 | Hypothetical protein | 228 | 2.00 | 8.99 | 5.18 |
| tetur03g10063 | Hypothetical protein | 294 | 7.02 | 3.40 | 5.15 |
| tetur04g05460 | 10-formyltetrahydrofolate dehydrogenase | 2727 | 1.39 | 1.78 | 2.27 |
| tetur05g01110 | ABC-transporter; class C (ABCC-16) | 4569 | 1.82 | 2.45 | 1.75 |
| tetur05g09325 | UDP-glucosyltransferase (UGT) | 1347 | 1.47 | 3.78 | 1.33 |
| tetur07g03960 | Apolipoprotein D | 603 | 1.70 | 1.80 | 1.42 |
| tetur07g03970 | Apolipoprotein D | 597 | 2.51 | 3.91 | 1.72 |
| tetur07g08269 | Hypothetical protein | 300 | 1.96 | 1.59 | 1.70 |
| tetur09g03890 | Hypothetical protein | 504 | 1.98 | 1.60 | 1.80 |
| tetur09g04920 | Apolipoprotein D precursor | 582 | 4.21 | 2.06 | 2.20 |
| tetur10g02090 | UDP-glycosyltransferase (UGT48) | 1305 | 2.02 | 1.45 | 1.68 |
| tetur10g02360 | Sialin | 1491 | 1.15 | 1.64 | 1.03 |
| tetur11g06390 | Hypothetical protein | 957 | 1.39 | 2.50 | 1.25 |
| tetur13g04230 | Hypothetical protein | 666 | 2.35 | 6.68 | 2.13 |
| tetur16g02380 | Carboxyl/cholin esterase (CCE40) | 1389 | 1.95 | 1.46 | 1.87 |
| tetur16g02420 | Carboxyl/cholin esterase (CCE43) | 744 | 1.92 | 2.51 | 1.29 |
| tetur17g03490 | Hypothetical protein | 243 | 1.43 | 1.84 | 1.17 |
| tetur18g01050 | Major facilitator superfamily domain | 1338 | 3.82 | 5.24 | 4.98 |
| **tetur20g01790** | **Intradiol ring-cleavage dioxygenase (DOG15)** | **885** | **4.30** | **4.98** | **3.60** |
| tetur21g00410 | Sodium-dependent glucose transporter 1 | 1377 | 1.40 | 2.79 | 1.10 |
| tetur24g01030 | Apolipoprotein D | 681 | 4.49 | 3.75 | 2.93 |
| tetur25g00650 | Cathepsin L (Pap42) | 987 | 2.87 | 2.19 | 2.12 |
| tetur29g00340 | Methyltransferase type 11 | 834 | 2.37 | 1.80 | 2.49 |
| tetur30g01770 | Hypothetical protein | 813 | 1.10 | 5.20 | 1.79 |
| tetur32g01960 | Carbonyl reductase (CBR1b) | 834 | 1.58 | 2.81 | 2.80 |
| tetur36g00190 | Hypothetical protein | 1389 | 1.48 | 3.83 | 2.65 |
| tetur86g00050 | Hypothetical protein | 597 | 1.89 | 5.42 | 1.32 |
| tetur92g00010 | Fibroin | 180 | 2.05 | 4.64 | 1.55 |

DOG15 is shown in bold.
