## Supplemental Table 3-2 for "Adaptation of an herbivorous arthropod to green tea plants by overcoming catechin defenses"

**Supplemental Table 3-2. List of 29 proteins highly expressed in tea-adapted KSM fed on tea leaves.** The levels of proteins were compared to tea-adapted KSM fed on bean leaves, non-adapted KSM fed on tea leaves, and TSSM fed on tea leaves.

| Database ID | Gene annotation | Molecular weight (kDa) | Log2 fold change | | |
|---|---|---|---|---|---|
| | | | vs Tea-adapted KSM on bean | vs TSSM on tea | vs Non-adapted KSM on tea |
| tetur01g00490 | Intradiol ring-cleavage dioxygenase (DOG1) | 28.8 | 3.39 | 3.04 | 1.57 |
| tetur01g03340 | tRNA/rRNA methyltransferase | 44 | 1.31 | 6.64 | 1.72 |
| tetur01g08680 | Carboxyl/cholinesterase (TuCCE01) | 45.7 | 2.81 | 3.22 | 1.31 |
| tetur02g05010 | UPF0692 protein C19orf54 homolog | 36.1 | 6.64 | 6.64 | 6.64 |
| tetur02g05890 | Tetratricopeptide repeat | 61.2 | 6.64 | 6.64 | 6.64 |
| tetur02g08310 | Dalao/Smarce1 related protein 2 (DalaoR2) | 63.1 | 6.64 | 6.64 | 6.64 |
| tetur03g07890 | Insulin-like growth factor binding protein | 101.6 | 6.64 | 3.57 | 2.46 |
| tetur03g08090 | Female lethal d homolog | 32.1 | 1.21 | 6.64 | 1.54 |
| tetur03g09470 | Alpha/beta hydrolase fold-5 | 32.4 | 6.64 | 6.64 | 6.64 |
| tetur03g91572 | Hypothetical protein | 29.9 | 5.49 | 6.64 | 6.64 |
| tetur04g04390 | S-Rnase | 26.4 | 1.34 | 2.19 | 2.32 |
| tetur04g04540 | Alpha/Beta hydrolase fold | 47.3 | 1.75 | 2.28 | 1.89 |
| tetur05g03950 | Nucleotide-sugar transporter-related | 42.2 | 6.64 | 6.64 | 6.64 |
| tetur06g00450 | Intradiol ring-cleavage dioxygenase (DOG4) | 29.4 | 1.27 | 2.55 | 1.9 |
| tetur07g05930 | Intradiol ring-cleavage dioxygenase (DOG7) | 30.1 | 1.94 | 6.64 | 2.03 |
| tetur07g07470 | ATG4 autophagy related 4 homolog A | 52.1 | 6.64 | 6.64 | 6.64 |
| tetur09g01660 | UDP-glucosyltransferase (UGT) | 52.4 | 2.44 | 2.75 | 1.19 |
| tetur14g02690 | Autophagy-related protein 12 | 12.4 | 6.64 | 6.64 | 6.64 |
| tetur14g03550 | 28S ribosomal protein S18c | 16.8 | 6.64 | 6.64 | 1.37 |
| tetur15g00180 | Hypothetical protein | 103.1 | 1.01 | 1.66 | 1.41 |
| tetur15g01540 | Cortactin-binding protein-2, N-terminal | 67.3 | 6.64 | 6.64 | 6.64 |
| tetur16g03290 | Hypothetical protein | 45.2 | 6.64 | 6.64 | 6.64 |
| **tetur20g01790** | **Intradiol ring-cleavage dioxygenase (DOG15)** | **32.5** | **2.22** | **1.81** | **3.56** |
| tetur23g00510 | Similar to Chaoptin precursor | 152.6 | 6.64 | 6.64 | 2.03 |
| tetur30g00040 | Hypothetical protein | 12.3 | 6.64 | 6.64 | 2.09 |
| tetur32g00060 | Similar to Glucosylceramidase precursor | 56.9 | 1.8 | 6.64 | 1.98 |
| tetur33g00190 | Srrm2 protein | 12.6 | 1.63 | 6.64 | 1.75 |
| tetur60g00020 | Legumain (TuLeg-9) | 49.4 | 2.88 | 6.64 | 1.18 |
| tetur143g00020 | Small Secretory Protein; Family F | 22.5 | 2.88 | 4.31 | 1.08 |

DOG15 is shown in bold.
