## Supplemental Table 4-1 for "Adaptation of an herbivorous arthropod to green tea plants by overcoming catechin defenses"

**Table S4-1**

**Supplemental Table 4-1. Specific activities of TkDOG15 and TuDOG15 toward catechins.** Enzymatic activity was measured with three independent experimental runs ($n$ = 3).

| Catechins | Specific activity (U/mg)[a] | | $p$-value[b] |
|---|---|---|---|
| | TkDOG15 | TuDOG15 | |
| EGCg | $4.0 \times 10^{-1} \pm 1.2 \times 10^{-2}$ | $2.4 \times 10^{-1} \pm 3.8 \times 10^{-2}$ | $0.002^{**}$ |
| EGC | $8.5 \times 10^{-1} \pm 5.2 \times 10^{-2}$ | $3.4 \times 10^{-1} \pm 1.1 \times 10^{-2}$ | $<0.001^{***}$ |
| ECg | $1.3 \pm 3.1 \times 10^{-2}$ | $1.0 \pm 8.1 \times 10^{-2}$ | $0.3^{NS}$ |
| EC | $6.0 \pm 2.4 \times 10^{-1}$ | $5.2 \pm 5.0 \times 10^{-2}$ | $0.004^{**}$ |

[a]Mean ± SD. [b]Student's $t$-test ($^{***}p < 0.001$, $^{**}p < 0.01$, $^{NS}p > 0.05$).
